# Evolutionary convergence and trophic diversity in hot vent and cold seep shrimps showcase a continuum of symbiosis

**DOI:** 10.1101/2025.08.27.672534

**Authors:** Pierre Methou, Margaux Mathieu-Resuge, Loïc N. Michel, Valérie Cueff-Gauchard, Hiromi Kayama Watanabe, Emily J. Cowell, Jonathan T. Copley, Roxanne Beinart, Magali Zbinden, Florence Pradillon, Marie-Anne Cambon, Chong Chen

## Abstract

Convergent evolution offers a powerful lens through which to examine the selective forces shaping life in extreme environments. In deep-sea hot vents and cold seeps, invertebrates have independently evolved symbioses with chemosynthetic bacteria, but repeated origins of such associations within a family remain rare. Here, we investigate the evolutionary emergence of chemosymbiosis in the shrimp family Alvinocarididae across 22 species collected globally. Electron microscopy identified a gradient of epibiotic bacterial colonization within the cephalothoracic cavity, ranging from absent to dense filamentous mats, suggesting distinct trophic strategies. Isotope and lipid trophic markers confirmed difference in reliance on chemosynthetic production among sympatric species with different bacterial colonization from a single vent. Phylogenetic analysis reveals at least two independent origins of chemosymbiosis, suggesting evolutionary convergence. Microhabitat association data further show that symbiotic phenotypes are most common in shrimps occupying the hottest, most geofluid-enriched microhabitats, though exceptions suggest contributions from additional ecological or physiological constraints. Our findings reveal many alvinocaridids as gradually evolving towards symbiosis, highlighting the importance of intermediate cases to understand the pathways to chemosymbiosis. This study contributes to a broader understanding of the predictability of evolutionary outcomes in distinct habitats such as vents, with broader implications for resilience of deep-sea ecosystems.

## Introduction

Repeated evolution of similar phenotypic traits within distinct lineages is often considered as the most compelling evidence for the action of natural selection [1,2]. The recurrence of similar adaptive solutions in response to comparable environmental contexts is unlikely to arise by chance alone [3,4]. This can result from parallel evolution towards a new phenotype if ancestral phenotypes are similar, or from convergent evolution when the ancestral phenotypes are distinct [4]. While evolutionary constraints can limit phenotypic variation, the repeated emergence of analogous traits act like replicated experiments of natural selection and shed light on the underlying factors and mechanisms that govern the adaptation of species [3].

At hydrothermal vents and other deep-sea chemosynthetic ecosystems such as hydrocarbon seeps, many invertebrate species have independently evolved a symbiotic partnership with chemosynthetic bacteria as an adaption to the particular conditions of these environments [5]. In the absence of sunlight, these associations result in ‘holobiont’ organisms where the host is dependent on the primary production of their microbial partners, which make use of chemical energy generated by geochemical gradients in the mixing zone where seawater collides with geofluids [6]. However, not all species there are symbiotrophic – several other feeding strategies such as bacterial grazing, deposit feeding, suspension feeding, scavenging and/or predation have been reported in other species endemic to chemosynthetic environments [7–9]. For most families with symbiotrophic members, the appearance of symbiosis is a unique event either at the common ancestor of the family such as siboglinid tubeworms, mussels in the subfamily Bathymodiolinae, paskentanid snails in the sister genera *Alviniconcha* and *Ifremeria* and yeti crabs in the family Kiwaidae [5,10–12], or in a single member of a family such as the squat lobster *Shinkaia crosnieri* within Munidopsidae [13]. A notable exception are peltospirid snails, in which chemosynthetic symbioses seem to have appeared at least twice, in two distantly related genera *Chrysomallon* and *Gigantopelta* [14], and potentially even a third case in *Peltospira gargantua* [15]. In the evolutionary history of these families, the acquisition of a chemosynthetic symbiosis is not necessarily concomitant with colonization of a chemosynthetic-based habitat, questioning the respective role of selective pressures from the environment, evolutionary constraints of these hosts, and historical contingencies in the making of such symbiotic relationships.

As with peltospirid snails, alvinocaridid shrimps likely acquired chemosymbiosis several times independently, once in the sister species pair *Rimicaris exoculata*/*R. kairei* and another in the species complex *R. chacei*/*R. hybisae* [16]. These shrimps host dense communities of filamentous bacteria in their cephalothoracic cavities and on the surface of their mouthparts that harbour a great metabolic versatility, adapted to different geochemical contexts depending on the hydrothermal fields they occupy [17–19]. Three of these shrimp species (*R. exoculata, R. hybisae* and *R. kairei*) display a hypertrophy of their symbiont-hosting surfaces, with an inflated cephalothoracic chamber and enlarged mouthparts [16], and are considered to have a nutrition based mainly on their symbionts [20,21] with a direct nutritional transfer demonstrated for *R. exoculata* [22]. These symbiont-hosting shrimps gather in dense swarms of thousands of individuals near the vent fluid orifices, often constituting the most dominant fauna there [23,24]. Conversely, several other species in the family or even genus such as *Nautilocaris saintlaurentae, Rimicaris acuminata*, and *Rimicaris variabilis* show limited colonisation by chemosymbiotic bacteria [25]. These species tend to feed on other food sources available such as bacterial mats and/or detritus [25–28]. In *R. chacei*, the presence of a dense bacterial colonization is not accompanied with the enlargement of cephalothorax, but results from isotopic and anatomical analyses indicate it partially depends on symbionts for nutrition in a mixed diet, while using other food sources [21,26,29]. This suggests that chemosymbiosis might be more widespread in alvinocaridid shrimps than previously identified simply by external morphology. Yet, the observation of bacterial colonization patterns remained limited to a few species and we still lack an overview of chemosymbiotic associations and diets in this family.

Our study aims to reconstruct the evolutionary history of chemosymbiosis in alvinocaridid shrimps and link it to the habitat they occupy. We address the following questions: 1) What are the patterns of bacterial colonization in different alvinocaridid species and how many time have dense bacterial colonization appeared? 2) Does alvinocaridid species with distinct colonization degrees exhibit different trophic strategies? 3) Is dense colonization by epibiotic filamentous bacteria linked to specific microhabitat occupancy along the environmental gradient of geofluid emissions in the local environment?

## Materials and Methods

### Sample collection

A total of 22 alvinocaridid shrimp species were gathered from samples originating from 32 different vent and seep localities around the globe (Table S1), collected during 31 different oceanographic expeditions between 2004 and 2023. They were mostly collected using suction samplers on either remotely operated vehicles (ROVs) or human-occupied vehicles (HOVs) and supplemented by samples from baited traps or epibenthic sledges (see Table S2 and S3 for detailed information). For scanning electron microscopy (SEM) observations, shrimp specimens were preserved either in 80% ethanol, frozen whole at −80°C, or fixed for 16h in 2.5% glutaraldehyde / filtered seawater solution, rinsed, and then preserved in filtered seawater solution at 4°C on-board of the ship. For specimens frozen whole on-board, post-fixation in 2.5% glutaraldehyde was done in the laboratory before SEM analyses. DNA extractions were carried out using individuals preserved in 80% ethanol or frozen at −80°C. Additional specimens collected at the Puy des Folles vent field on the Mid-Atlantic Ridge (MAR) during the BICOSE 3 expedition were stored frozen at −80°C for stable isotopes and lipid analyses (Table S4 and S5).

### Stable isotope and lipid analysis

The Puy des Folles hydrothermal site was selected to investigate stable isotope and fatty acid profiles of alvinocaridids because it offered the widest range of species showing distinct bacterial colonization patterns available to us, with four different species within a single hydrothermal vent locality including *Rimicaris exoculata, R. chacei, Mirocaris fortunata*, and *Alvinocaris markensis*. For stable isotopes analyses, pieces of abdominal muscle and mouthparts (scaphognathites – i.e., exopodite of the second maxilla – and exopodites of the first maxillipeds, hereafter called just exopodites)from three specimens per species were isolated, freeze dried and homogenized with a manual potter. Homogenized pieces of abdominal muscles from the same individuals were also used for fatty acid analyses.

Measurements of stable isotope ratio were performed by continuous flow–elemental analysis isotope ratio mass spectrometry (CF-EA-IRMS) at University of Liège (Belgium), using a vario MICRO cube C-N-S elemental analyser (Elementar Analysensysteme GMBH, Hanau, Germany) coupled to an IsoPrime100 isotope ratio mass spectrometer (Isoprime, Cheadle, United Kingdom). Isotopic ratios were expressed in ‰ using the widespread δ notation [30] relative to the international references: Vienna Pee Dee Belemnite (for carbon), atmospheric air (for nitrogen) and Vienna Canyon Diablo Troilite (for sulphur). IAEA (International Atomic Energy Agency, Vienna, Austria) certified reference materials sucrose (IAEA-C-6; δ^13^C = −10.8 ± 0.5‰; mean ± SD), ammonium sulphate (IAEA-N-2; δ^15^N = 20.41 ± 0.12‰; mean ± SD), and barium sulphate (IAEA-SO5, δ^34^S = 0.5 ± 0.2‰) were used as primary analytical standards. Sulfanilic acid (Sigma-Aldrich; δ^13^C = −25.6 ± 0.4‰; δ^15^N = −0.13 ± 0.4‰; δ^34^S = 5.9 ± 0.5‰; means ± SD) was used as a secondary analytical standard. Standard deviations on multi-batch replicate measurements of secondary and internal lab standards (seabass muscle) interspersed with samples (one replicate of each standard every 15 analyses) were 0.2‰ for δ^13^C, 0.1‰ for δ15N and 0.3‰ for δ^34^S. Δ^13^C was defined as the δ^13^C isotopic ratio of abdominal muscle tissue (the “consumer”) from which was subtracted the δ^13^C isotopic ratio of mouthparts more or less colonized by symbiotic bacteria (the “nutrition source”) depending on the shrimp species. Following the same principle, Δ^15^N and Δ^34^S were calculated for each individual as follows: Δ^15^N = δ^15^N _muscle_ - δ^15^N_mouthpart_and Δ^34^S = δ^34^S_muscle_ - δ^34^S_mouthpart_.

A minimum of 2 mg of abdominal tissue (n = 3 per species) powder were placed in pre-combusted glass vials and 6 ml of solvent mixture (CHCl3: MeOH, 2:1, v:v) were added to extract lipids. Lipid extracts were then flushed under nitrogen, sonicated, vortexed, and rested at −20°C for 24h. As described in a previous study [31], an aliquot of total lipid extract was used for transesterification. Briefly, after adding C23:0 as internal standard (free fatty acid form), lipids were first transesterified with 1ml KOH-MeOH and then with 1.6ml of H_2_SO_4_-MeOH, to form fatty acid methyl ester (FAME) that were then rinsed with distilled water saturated in hexane, and recovered within 0.4ml of hexane. FAME were analyzed on a TRACE 1300 gas chromatograph (Thermo Scientific) programmed in temperature and equipped with a splitless injector, a ZB-WAX column (30m×0.25mm IDx0.2μm) and a flame-ionisation detector, using hydrogen as vector gas. Obtained chromatograms were processed with Chromelon 7.2 (Thermo Scientific). Sixty-four FAME were identified by comparing their retention time with references from commercial mixtures (37 components FAME, PUFA1 and PUFA3, Sigma) and from house-made standards GC-MS certified; as well as by checking certain samples using mass spectrometry (GC-MS TRACE 1300 coupled to an ISQ 7000 mass spectrometer, equipped with the same apolar column and using the same program as GC-FID). Quantification of FAME in µg was based on the internal standard recovery, and then expressed in mg g^−1^ of dry weight. Fatty acids relative proportions were expressed as mass percentages (%) of the total identified fatty acids.

All statistical analyses with Kruskal-Wallis and Wilcoxon tests were performed in a R v. 4.4.1 statistical environment [32].

### Scanning electron microscopy

SEM observations were carried out on 17 alvinocaridid species sampled to investigate the bacterial colonization on the cephalothoracic cavity. 48 dissected branchiostegites (inner side of the cephalothoracic cavity) and mouthparts (scaphognathites and exopodites) of individuals fixed in 2.5% glutaraldehyde were dehydrated with an ethanol series from 50% to 100% ethanol by 10% increments or directly into 100% ethanol for individuals preserved in 80% ethanol. Samples were then dehydrated for 5 h in a critical point dryer CPD 020 (Balzers Union, Balzers, Liechtenstein) and gold-coated with a SCD 040 sputter-coater (Balzers Union). Observations and imaging were performed using a Quanta 200 SEM (FEI-Thermo Fisher, Hillsboro, OR, USA).

We defined a colonization score for consistency when comparing our multiple SEM observations. Branchiostegites (Br) were divided in three equivalent areas (anterior, center and posterior) and mouthparts (Mp) in three features: the central surface, the marginal setae on the edge of the mouthparts, and the additional bacteriophore setae covering the surface, present only in some species (Figure S1). For each area/feature, a score from 0 to 4 was attributed according to the following criteria, 0: no visible bacterial colonization; 1: a single layer of rod-shaped bacterial mats; 2: colonization by rod-shaped bacteria and sparsely distributed filamentous bacteria; 3: localized spots colonized by filamentous bacteria; 4: extremely dense colonization by filamentous bacteria, the surface of shrimp cuticle barely visible. The final colonization score is obtained by the addition of each areas/features colonization scores for a total score of 0 to 24. We reanalysed the SEM dataset from a previous study [25] to define the colonization scores of species analyzed by this study (*R. variabilis, R. acuminata* and *N. saintlaurentae*). We based our colonization scores of *R. exoculata* and *R. kairei* from descriptions of their cephalothoracic symbioses in the literature [17,19].

### DNA extraction and sequencing

We performed DNA extractions on individuals from the same 22 alvinocaridid species sampled using pieces of abdominal tissues or the 4th/5th pleopods with the DNeasy Blood & Tissue kit (Qiagen, Hilden, Germany) following the manufacturer’s instructions. Partial fragments of the nuclear 18S rRNA (905 bp), and the mitochondrial 16S rRNA (515 bp) and the cytochrome oxidase *c* subunit I (COI) gene (743 bp) were amplified respectively with the 18S-1f & 18S-5r, the Cari16S-1F and Cari16S-1R [33], and the CariCOI-1F and CariCOI-1R primers [21]. We used ExTaq polymerase (Takara) or GoTaq polymerase (Promega) in 25μl reactions for PCR amplifications with the following cycling conditions: 40 cycles at 51°C as the annealing temperature for 18S, 35 cycles at 55°C for 16S, and 35 cycles at 50°C for COI. Bi-directional Sanger sequencing was conducted by Eurofins (Köln, Germany) or by the FASMAC Corporation (Kanagawa, Japan) for the different PCR products. We used Geneious Prime® 2023.1.2 (https://www.geneious.com) to edit and align sequence chromatograms. We carried out phylogenetic reconstruction using the Mr Bayes plugin v3.2.6 [34] in Geneious. Bayesian inference was run with the GTR + G + I substitution model and 1 million MCM steps, discarding the first 10 000 steps and with a subsampling frequency of 4000 steps. Best fitting model was selected with Modeltest [35] within Geneious. Ancestral state reconstruction was conducted with the ace function of the phytools v2.4-4 package [36] in RStudio v4.4.1.

## Results

### Stable isotope and lipid analysis

Within a single hydrothermal vent field - Puy des Folles on the Mid Atlantic Ridge - each of the four alvinocaridid species displayed clearly distinct fatty acid and stable isotope composition (Figure 1). The δ^15^N of most alvinocaridid mouthparts was significantly lower than the δ^15^N of abdominal muscles from the same species (Wilcoxon test; W= 118, p-value < 0.01), except for *A. markensis* that showed the opposite trend with equivalent or higher δ^15^N values of mouthparts compared to abdominal muscles. Inter-specific differences in nitrogen trophic shift (Δ^15^N) were present (Kruskal–Wallis; χ^2^ = 9.46, p-value < 0.05), with higher values for the two *Rimicaris* species (*R. exoculata*: Δ^15^N = 3.17 ± 0.17; *R. chacei*: Δ^15^N = 3.34 ± 0.95) compared to *A. markensis* (Δ^15^N = −0.05 ± 1.53). The δ^13^C did not vary between muscles and mouthparts (Wilcoxon test; W= 90, p-value > 0.05), but varied among species (Kruskal–Wallis; χ^2^ = 17.79, p-value < 0.001) with a significant ^13^C-enrichment for *R. exoculata* and *M. fortunata* compared to *R. chacei* and *A. markensis*. Carbon trophic shift (Δ^13^C) was high in *R. chacei* compared to other alvinocaridids, which exhibited low Δ^13^C values in *R. exoculata* and *M. fortunata* or even negative values for *A. markensis*. As for carbon, δ^34^S did not vary between tissues (Wilcoxon test; W= 57, p-value > 0.05) but varied among species (Kruskal–Wallis; χ^2^ = 15.9, p-value < 0.001) with a significant ^34^S-depletion for *M. fortunata* and *A. markensis* compared to the two *Rimicaris* species (Figure S2).

**Figure 1.**
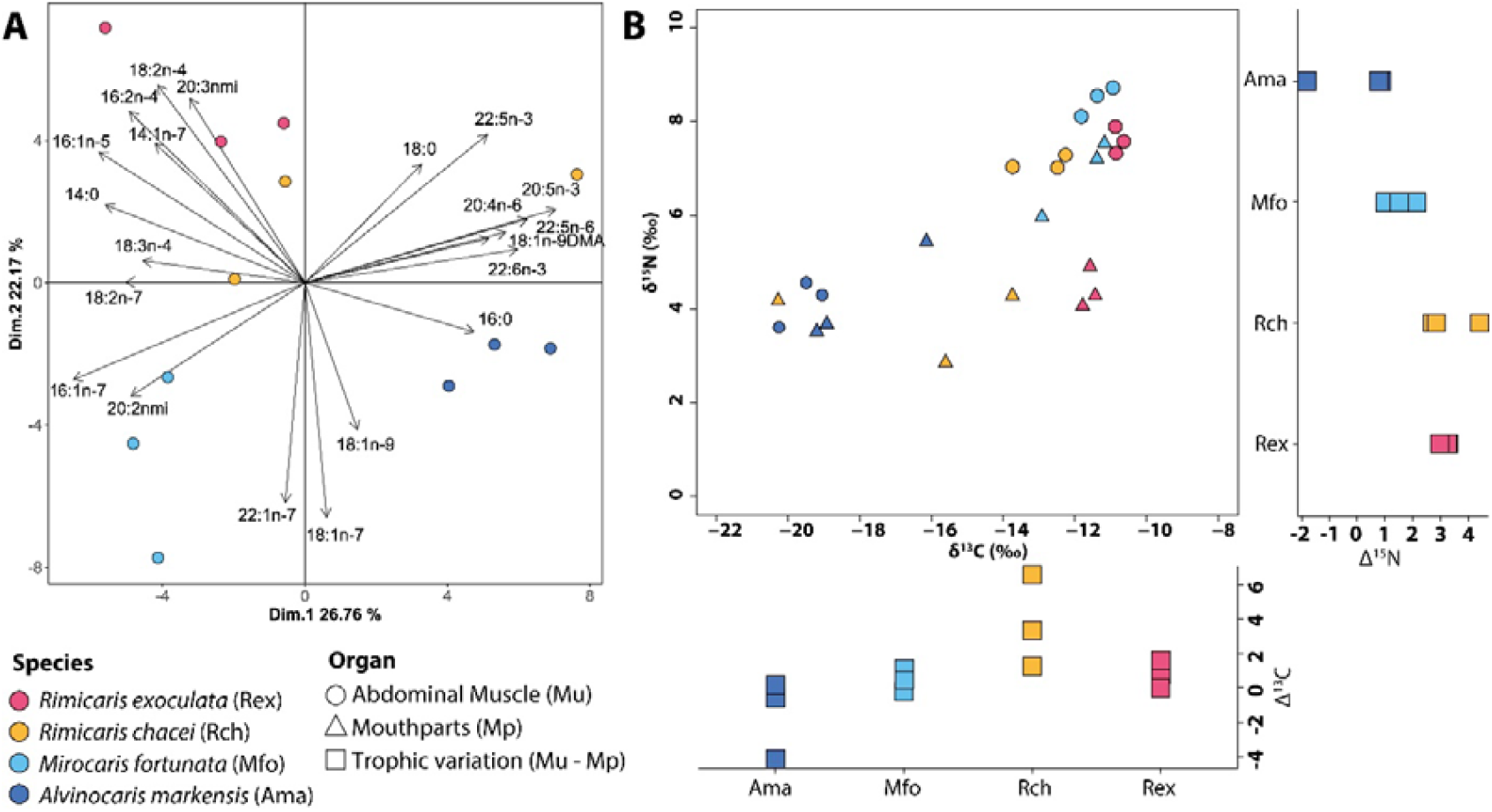
Trophic strategies of alvinocaridid species displaying different degree of bacterial colonization at the Puy des Folles vent field. Colors indicates shrimp species and shape indicates organ analyzed. **A.** Principal Component Analyses (PCA) of fatty acid composition (expressed in mass %) of the abdominal muscles from the distinct alvinocaridid species **B**. Carbon and nitrogen isotopic ratios of abdominal muscles and mouthparts from alvinocaridid shrimps. Δ^13^C and Δ^15^N display variations of carbon and nitrogen ratios between muscles and mouthpart tissues of each individual.

Overall, species differed in their total fatty acid content (Kruskal–Wallis test, H25 = 9.67, p-value < 0.05; Table S5). *Rimicaris* species had the higher significant total contents with 207.0 ± 34.0 and 181.0 ± 27.0 mg g^−1^ for *R. exoculata* and *R. chacei*, respectively. *M. fortunata* (24.7 ± 2.4 mg g^−1^) and *A. markensis* (20.8 ± 0.9 mg g^−1^) contained fatty acid contents 10 and 8.3 times lower than *R. exoculata*, respectively. The fatty acid compositions of shrimp abdominal muscles varied significantly among species (PERMANOVA, df = 3, F = 5.34, r^2^ = 0.59, p-value < 0.001), as highlighted by the PCA on which the two first axis explained 48% of the total inertia (26.76 % on the first axis and 22.17 % on the second, Figure 1A). The fatty acids profiles of the two *Rimicaris* species were positively related to axis 2, while that the two other species were negatively related to this axis. Individuals of *R. exoculata* and two of *R. chacei* were characterized by high proportions of short chain fatty acids mainly composed of 14:0, 14:1n-7, 16:1n-5, 16:2n-4, 18:2n-4, 18:3n-4, 18:2n-7 and by the 20:3nmi (non-methylene interrupted) (Figure 1A). Individuals of *A. markensis* and one of *R. chacei* were discriminated positively on the first principal axis and characterized by higher proportions of longer chain fatty acids, such as 20:4n-6, 20:5n-3, 22:5n-3, 22:6n-3 and the 18:1n-9DMA (dimethylacetal) (Figure 1A). Individuals of *M. fortunata* were negatively discriminated on the first and second principal axis, and characterized by higher proportions of monounsaturated fatty acids, such as 16:1n-7, 18:1n-7, 18:1n-9, 22:1n-7 and the 20:2nmi (Figure 1A).

### SEM observations

For most of the 22 alvinocaridid species observed, some general trends of bacterial colonization on branchiostegites and on mouthparts emerge (Figure 2 and Table S2). Overall, branchiostegites were generally devoid of colonization with only a few spots of rod-shaped bacteria in a single layer or a few scattered filamentous bacteria, most often on the anterior one-third (Figure 2; exemplified by *R. leurokolos* and *M. fortunata*). Mouthparts – scaphognathites and exopodites – show similar colonization patterns on most of their surfaces but contrast by a denser colonization on the plumose setae structures aligned along their margins (Figure 2; exemplified by *R. leurokolos* and *M. fortunata*). This results in total colonization scores between 3 and 10 score for most alvinocaridid species.

**Figure 2.**
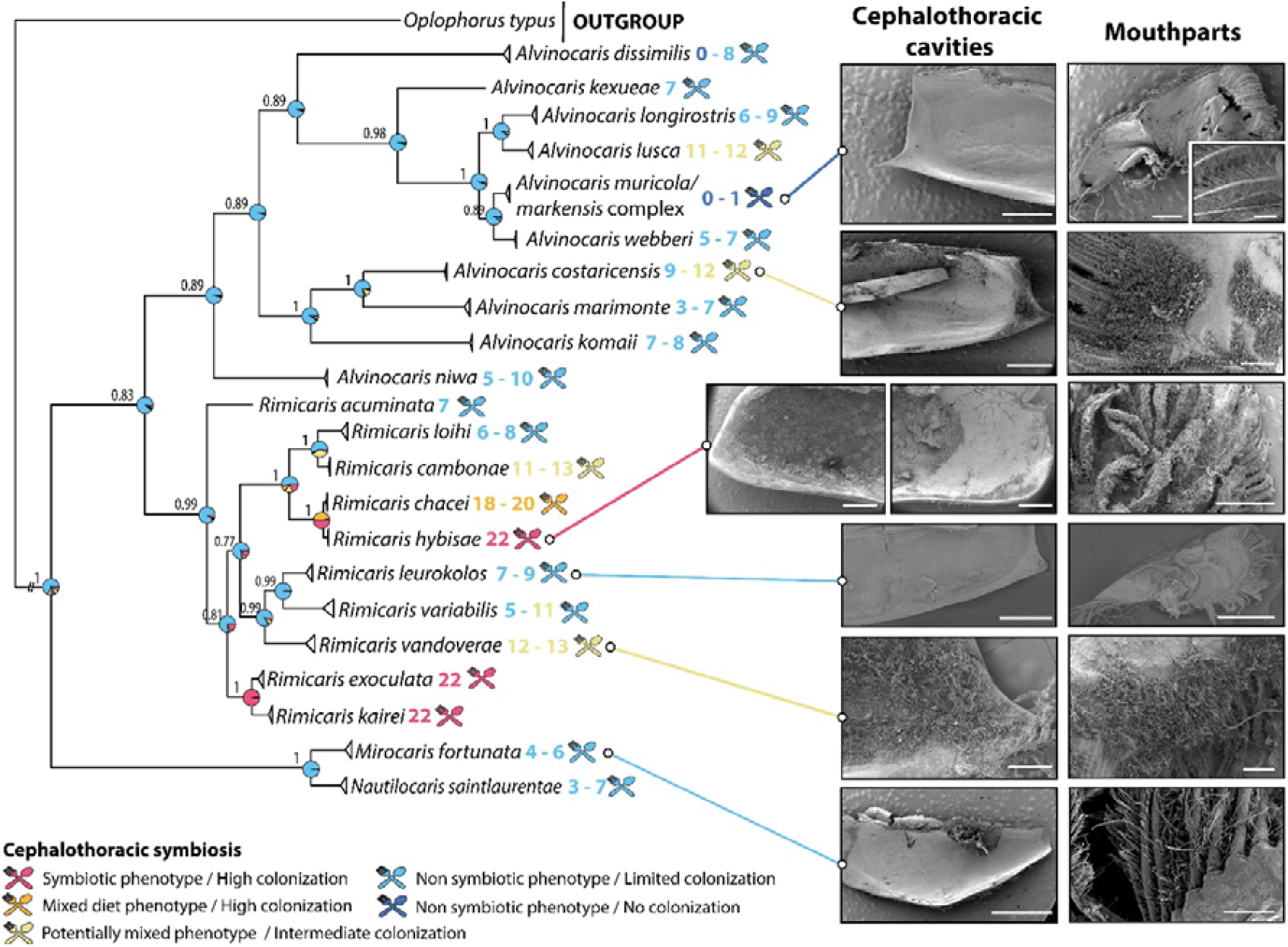
Evolutionary history of the chemosynthetic symbiosis in alvinocaridid shrimps **(left)** Phylogenetic tree of Alvinocarididae based on Bayesian inference (GTR + G + I model) generated from a 2177 bp concatenated alignment comprising partial fragments of the 18S, 16S, and COI genes. Coloured numbers associated to each species indicate the range of colonization scores defined from SEM observations and colours correspond to the different categories of chemosymbiotic phenotypes (see legend). Numbers on each node indicate Bayesian Posterior Probabilities (BPP) of the phylogeny and pie charts represent ancestral phenotypes displaying posterior probabilities of each node being in each category of phenotypes. **(right)** Representative examples of SEM observations for the cephalothoracic cavities and mouthparts linked to the alvinocaridid species observed. Scale bars from top to bottom for cephalothoracic cavities: *A. muricola* overview (1 mm); *A. costaricensis* (2 mm); *R. hybisae* (2 mm); *R. leurokolos* (1 mm); *R. vandoverae* (500 μm); *M. fortunata* (2.5 mm). Scale bars from top to bottom for mouthparts: *A. muricola* overview (1 mm) and close up (100 μm); *A. costaricensis* (250 μm); *R. hybisae* (1 mm); *R. leurokolos* (1 mm); *R. vandoverae* (250 μm); *M. fortunata* (50 μm). Additional SEM observations for all studied species are provided in Figure S4 and S5.

Apart from these general trends, some species show more specific colonization patterns (Figure 2 and Table S1). Among them, *Rimicaris hybisae* and *R. chacei* differ drastically from the other species by a very dense colonization of filamentous bacteria on all structures except on the posterior part of the branchiostegite. These two species were mainly distinguished by a more advanced colonization of dense filamentous bacteria covering the anterior two-thirds of the branchiostegite for *R. hybisae* (Figure 2) and the anterior one-third only for *R. chacei*. These two species also show additional bacteriophore setae structures (less numerous in *R. chacei*) on the entire mouthpart surfaces offering additional colonisation areas compared to other alvinocaridids, further enhancing their colonization score (*R. hybisae*: 22; *R. chacei*: 18 to 20). Based on the literature, a similarly dense colonization of filamentous bacteria on all structures except on the posterior third of the branchiostegite with a colonization score of 22 can be attributed to *R. kairei* and *R. exoculata* [17,19].

Two other *Rimicaris* species – *R. vandoverae* and *R. cambonae* – also departs from the general pattern with a dense colonization by filamentous bacteria on the anterior one-third of their branchiostegite (Figure 2), and to some extent, on their mouthpart surfaces as well, leading to total colonization scores of 12 to 13 for *R. vandoverae* and of 11 to 13 for *R. cambonae*. In addition, two *Alvinocaris* species – *A. lusca* and *A. costaricensis* – consistently exhibit dense spots of filamentous bacteria, not only on the marginal setae but also on the surfaces of their mouthparts in all individuals observed (Figure 2), as well as on the anterior one-third of the branchiostegites in some individuals (*A. lusca* scores: 11 to 12; *A. costaricensis* scores: 9 to 12). Finally, in contrast to all other species, *A. markensis* shows a nearly complete absence of bacterial colonization on all of these structures (Figure 2), even on marginal setae of their mouthparts which were typically colonized in all other species (Figure 2), leading to a score of 0 or 1.

### Phylogenetic and Ancestral state reconstruction

Our phylogenetic reconstruction of Alvinocarididae based on multi-gene alignment (*COI, 16S* and *18S*) using Bayesian inference (Figure 2) recovered a strongly supported monophyly of the family (Bayesian Posterior Probability (BPP) = 1) as well as the monophyly of the genera *Rimicaris* (BPP = 0.99) and *Alvinocaris* (BPP = 0.89). The clade formed by *Mirocaris fortunata* + *Nautilocaris saintlaurentae* was the early-diverging lineage, sister to all the other alvinocaridid shrimps. All nodes were well-supported (BBP > 0.8), except one node within *Rimicaris* (BPP = 0.77). Ancestral state reconstruction supported two distinct origins for the appearance of a symbiotic phenotype: once at the common ancestor of *R. exoculata* and *R. kairei* and once in the *R. chacei*/*R. hybisae* species complex (Figure 2). It also supported multiple origins for the appearance of mixed or potentially mixed phenotypes (Figure 2). Ancestral state reconstruction of habitats revealed that a vent origin was most likely for the most recent common ancestor of Alvinocarididae with a secondary colonisation of hydrocarbon seeps in *A. costaricensis* and in the common ancestor of one of the two major clades within genus *Alvinocaris* clade (Figure S3).

### Habitat and colonization score

The relationship between the colonization score of an individual and its type of habitat was not entirely straightforward (Figure 3). All species with high colonization scores – i.e., score > 18 – such as the *R. exoculata, R. kairei, R. hybisae/R. chacei* complex – were always found in close proximity to vent fluid emissions. However, not all alvinocaridid species occupying similar habitat, such as *R. leurokolos* or some individuals of *R. variabilis*, necessarily showed such high colonization scores (Figure 3). Indeed, *R. leurokolos* shrimps showed more similar colonization scores to *A. longirostris* or *A. dissimilis* individuals from the same site but inhabiting cooler habitats within mussel beds or near *Shinkaia* aggregates. In addition, species with mid-range colonization score – i.e., score >10 – like *R. vandoverae, R. cambonae*, or *A. lusca* were found to inhabits cooler habitats than chimney surfaces such as *Alviniconcha* snail colonies or even cooler ones such as mussel beds. *Alvinocaris markensis* which displayed lowest colonization score were always found in peripheral areas away from the venting fluid.

**Figure 3.**
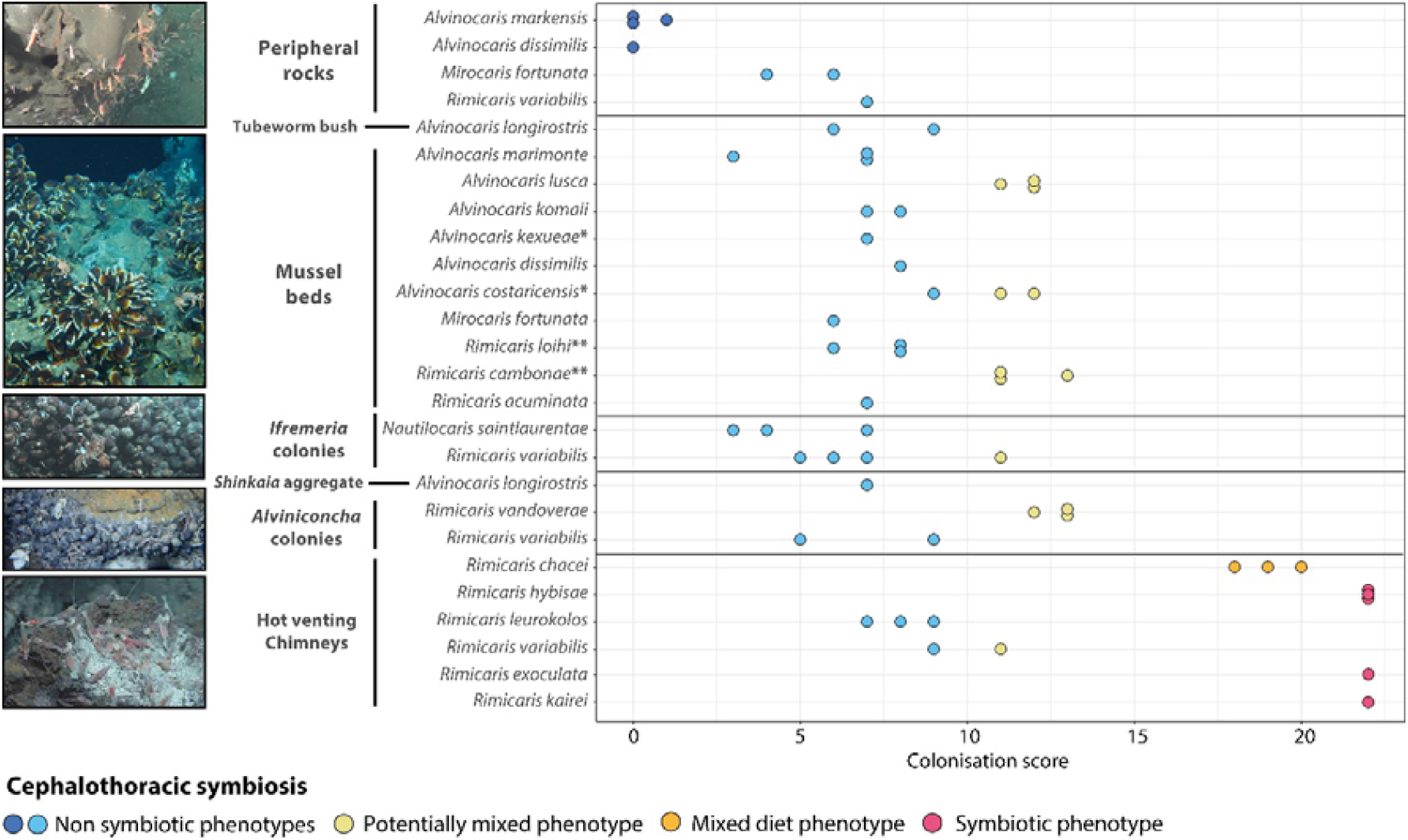
Individual colonization scores of each alvinocaridid species as a function of their habitat along the geochemical gradient. The habitats are ordered according to their distance to the venting or seepage emissions from close distance on the bottom and further away on top of the plot. Sampling of Alvinocaridids shrimps off New Zealand with an epibenthic sledge did not allow to associate a habitat type with their individual colonization scores. *: Species collected from methane seeps. **: *R. cambonae* and *R. loihi* were collected within a mussel bed in close vicinity to a hot venting habitat. These shrimps were found in both habitats [46] with individuals likely moving from one to another. Habitats types of alvinocaridid shrimps are illustrated by (from top to bottom): a peripheral rock habitat at Puy des Folles (Mid Atlantic Ridge); a mussel bed habitat at Minami-Ensei (Okinawa Trough); a *Ifremeria* snail colony at Susu Knolls (Manus Basin); an aggregation of *Alviniconcha* snails at Snail site (Mariana back-arc basin); a shrimp assemblage near hot venting fluids at PACMANUS (Manus Basin).

## Discussion

This study shows the presence of distinct trophic strategies among species of alvinocaridid shrimps that display different degrees of bacterial colonization within the same locality, as supported by our stable isotope and fatty acid analyses (Figure 1 & S1). As indicated by previous studies, the nutrition of *Rimicaris exoculata* is mainly based on its bacterial ectosymbionts [17,21,22]. Here, we recorded important values for Δ^15^N between the host organs (i.e., mouthparts) and muscle tissues of this species. This is generally expected between a consumer and its food items, which consistent with ectosymbionts being a major food source for this species. This is further supported by a fatty acid composition enriched in 16:2n-4 and 18:2n-4 sulfuric oxidative filamentous bacterial markers [37]. On the other end of the spectrum, *Alvinocaris markensis* showed a near absence of colonization, which is consistent with a non-symbiotic nutrition. The isotopic composition of its mouthparts and muscles were similar (absence of marked trophic shift), which is expected for two tissues synthesized independently by the same animal and does not suggest any trophic relationship. Their fatty acid compositions were also enriched in omega-3 and -6 long chain polyunsaturated fatty acids, which can be either associated to the ingestion of phytodetritus [37] and/or biosynthesised by bacteria [38,39]. In the same way, the limited bacterial colonization observed in *M. fortunata* is in line with limited inputs from its symbionts, demonstrated by a lower trophic shift and a distinct monounsaturated fatty acid composition often associated to thio-oxidizing bacteria [40]. Our trophic marker data on *R. chacei* individuals are also consistent with previous works suggesting a mixotrophic diet for this species [21,29].

Taken together, our results illustrate niche theory predicting that sympatric species are partitioned in their feeding habits and resource use to avoid competitive exclusion among the four studied species [41]. Through their symbiotic associations, host species such as *Rimicaris* are likely to access novel niches that are expanded by the addition of symbiont’s physiological and metabolic capacities [5]. A partnership with symbionts likely guarantees access to larger and more consistent quantities of food, as highlighted by the higher total fatty acid contents found in these two species. In chemosynthesis-based ecosystems, niche partitioning in relation to symbioses have been demonstrated in *Alviniconcha* snails where different hosts harbour variable symbiont phylotypes within the same region [42]. In alvinocaridid shrimps, the partitioning among sympatric species seems to be driven by various degrees of reliance on the production of their filamentous bacterial symbionts, from none to full dependence.

By expanding our assessment of bacterial colonization patterns to a global dataset of 22 alvinocaridid species, we revealed the presence of a gradient of colonization phenotypes within the family, with lineages associated with symbiotrophic, mixotrophic (or potentially mixotrophic) diets. This suggests an evolutionary convergence with multiple emergences of symbiotic phenotypes along the evolutionary history of Alvinocarididae (Figure 2), implying that this group has high flexibility in the development of distinct trophic strategies. Although uncommon in other chemosymbiotic holobionts, nutritional flexibility with limited contribution from heterotrophy has also been shown in several bathymodioline mussels which retain some ability to filter-feed [43,44], or *Alviniconcha* snails which can still digest some food with their reduced gut [45]. In alvinocaridid shrimps, this trophic flexibility can be further extended when considering the case of the *R. chacei*/*R. hybisae* species complex that display a developmental plasticity during their metamorphosis in the colonization surfaces of their symbiont-hosting organs, allowing individuals to perform either mixotrophic or symbiotrophic feeding depending on the environment [16]. This developmental variability could be due to competitive interactions for niche occupancy with *R. exoculata* at juvenile stages promoting a shift toward mixotrophy on MAR vents [24], while *R. hybisae* in the Mid-Cayman Spreading Centre has no local competitor and can fully develop its symbiotrophic phenotypes [23]. We suggest that other cases of niche partitioning are driving feeding strategies and symbiont colonization in alvinocaridids such as between *A. marimonte, R. cambonae*, and *R. loihi* living in sympatry on the Mariana Arc [27,46].

Our analyses also support habitat specialization as a key factor shaping symbiotic relationships in alvinocaridid shrimps (Figure 3). At an ecosystem-scale, the acquisition of sulfur-oxidizing or methanotrophic symbionts in bathymodioline mussels and vesicomyid clams coincides with their repeated colonisations in different types of chemosynthetic habitats along their evolutionary histories [11,47]. Similarly, the origin of chemosymbiosis in the abyssochrysoidean snails coincide with the Aptian oceanic anoxic event which allowed them to colonize hydrothermal vents, though the other major clade that split at the same time, containing genera such as *Provanna* and *Desbruyeresia*, did not evolve symbiosis [48]. Symbiosis in ‘vestimentiferan’ worms in the family Siboglinidae also evolved only once at the start of their association with reducing habitats, but its origin traces back widespread reducing, anoxic sediments instead of sparse and ephemeral vents or seeps [49]. These scenarios contrast with the emergence of chemosynthetic symbiosis in alvinocaridid shrimps, which is not directly linked to their colonization of reducing habitats, as our ancient habitat reconstruction indicated that association with hydrothermal vents is ancestral to the family. Instead, the symbiosis appeared secondarily, more similar to the case of some peltospirid snails [14,15]. Yet, at the microhabitat scale, the habitat preference of alvinocaridid species along the geochemical gradient appears to be important, as all species with high colonization scores occupy the hottest parts of the vent chimney with the greatest exposure to high concentrations of geofluids. This emphasizes the strong influence of the geochemical gradient in structuring the distribution and zonation of chemosynthetic holobiont species [5,6]. However, this link is not always clear-cut, with some alvinocaridid species of potentially mixed phenotypes inhabiting coolers habitats (e.g., *A. lusca* and *R. cambonae*) and species displaying limited epibiont colonization inhabiting hot venting areas (e.g., *R. leurokolos*), suggesting that other factors, including the co-occurrence with other abundant holobionts, are also at play for establishment of their realized niches.

The limited number of known chemosynthetic symbioses in crustaceans compared to annelids and molluscs [5,50] may be linked to evolutionary constraints of the phylum such as their moulting cycle. In symbiotic insects, moulting only during the early phases of their life cycles poses a challenge to the transmission and maintenance of symbiotic partnerships [51], which is further enhanced in crustaceans that moult throughout their adult life. In *R. exoculata*, the moulting does not hamper the establishment of a chemosynthetic symbiosis but the very rapid moulting frequency, estimated at every ten days, imposes a rapid renewal rate of the symbiont community [52], probably resulting in a significant metabolic cost. The high mobility of decapod crustaceans could also be another barrier to the emergence of symbiotic relationship in this group. Indeed, global functional trait analyses of hydrothermal vent ecosystems have highlighted that most species with a symbiotic feeding mode were sessile or only slightly mobile [53]. A high motility allows species to significantly expand accessible food resources instead of relying on a single trophic strategy, which could explain the gradual variations of symbiont colonization phenotypes we observe among alvinocaridid species.

These two factors – i.e., moulting and mobility – should also be mentioned as potential sources of bias in our observations which could contribute partly to the observed inter-individual variability of bacterial colonization observed. For our observations, we excluded individuals just after moulting – characterised by soft carapaces and a lack of mineral deposition – that have reset their epibiont colonization [52]. However, we cannot rule out that different individuals observed could be at slightly different moult – and thus bacterial colonization – stages. Similarly, with alvinocaridid shrimps being mobile, we cannot be sure if the sampling habitat differs or not from the habitat of that individual in the past few hours/days, especially for species known to occupy a wide range of habitat types such as *A. longirostris, R. variabilis* or *R. chacei* [24,28] (see also Figure 3). However, these potential biases are unlikely to affect the general trends we highlight, with several species, such as *R. leurokolos*, being restricted to a single type of habitat [28,54].

Overall, rather than a division in two categories of symbiotic and non-symbiotic species, our results indicate that chemosymbiosis gradually emerged in alvinocaridid shrimps, under the influence of the geochemical gradient but not only that. Our observations highlight a more complex and ongoing dynamic in the emergence of symbioses in vent and seep endemic species. As *R. chacei* and other shrimps with mixotrophic diet, several species in other phyla could be partially symbiotic. To some extent, mixotrophy has already been demonstrated in bathymodioline mussels. Many other examples inferred from stable isotopic compositions and/or observations of bacterial ectosymbionts could represent cases of partially chemosymbiotic species such as the amphipod *Exitomelita sigynae* [55] or the small vent snails *Peltospira smaragdina* and *Cyathermia naticoides* [8,56]. We argue that studying these gradual cases of chemosymbiosis is as essential as studies of symbioses from emblematic foundational species to fully understand the evolutionary dynamics of how symbiotic relationships are established.

Our results also have significant implications for the conservation of hydrothermal vent ecosystems. A recent study [53] highlighted the high proportion of functionally unique species with little redundancy in hydrothermal vents resulting in rapid collapse of these ecosystems using simulated species extinction – but the feeding modes of all alvinocaridids were categorized as bacterivores. Our work suggests there is more nuance with a continuum in terms of feeding modes within this family. Rather than diminishing the conclusions of that study [53], we further enhance the view of high functional uniqueness in vent endemic species which points again to the vulnerability of these environments in the face of mining threats.

## Conclusion

By investigating bacterial colonisation and trophic markers in alvinocaridid shrimps in an unprecedented global-scale study of 22 species, we underscore that the evolution of chemosynthetic symbiosis in alvinocaridid shrimps is a dynamic and recurrent process shaped by complex factors including ecological context, developmental plasticity, and phylogenetic constraints. The independent emergence of densely colonized symbiotic phenotypes, coupled with a spectrum of intermediate forms, points to evolutionary convergence driven by strong environmental selection. Yet, the incomplete correspondence between symbiosis and habitat, and the persistence of mixed trophic strategies, highlight the role of other factors like the physiological and developmental constraints due to moulting and mobility in modulating evolutionary trajectories. Our study exemplifies how complex traits such as symbiosis at vents and seeps can, and are being, evolved incrementally, through flexible and sometimes facultative intermediates, rather than through abrupt transitions as is seen in other groups like abyssochrysoid snails and vesicomyid clams. This continuum challenges simplistic categorizations of broader taxonomic groups inhabiting vents and seeps under one or a couple of feeding modes, since Alvinocarididae clearly includes a whole spectrum of feeding modes. Our work contributes to a more nuanced understanding of the repeatability — and limitations — of evolution in ‘extreme’ environments.

## Acknowledgments

We thank the captain, crew, and scientists on-board R/V *Pourquoi Pas?* during research expeditions BICOSE2014 (https://doi.org/10.17600/14000100), BICOSE2 (https://doi.org/10.17600/18000004), BICOSE3 (https://doi.org/10.17600/18002399), BIOBAZ (https://doi.org/10.17600/13030030), HERMINE2 (https://doi.org/10.17600/18001851), and WACS (https://doi.org/10.17600/11030010), R/V *Atalante* during expeditions CHUBACARC (https://doi.org/10.17600/18001111), FUTUNA3 (https://doi.org/10.17600/12010040), and MOMARSAT21 (https://doi.org/10.17600/18001296), R/V *Shinsei Maru* during expeditions KS-21-20 and KS-22-2, R/V *Natsushima* during expedition NT15-02, R/V *Kaimei* during expedition KM23-E05, R/V *Kaiyo* during expedition KY14-01, R/V *Yokosuka* during expeditions YK10-11, YK19-10, YK16-E02, YK22-05, YK23-06, and YK23-16, R/V *Haiyang 6* during expedition HYDZ6-2020-05, R/V *Tangaroa* during expeditions TAN0411, TAN1206, and TAN2102, R/V *Ka’imikai-o-Kanaloa* during expeditions KOK0506 and KOK0507, R/V *James Cook* during expedition JC82, R/V *Falkor (too)* during expedition FKt231024, R/V *Atlantis* during expedition AT42-03 and R/V *Thomas G. Thompson* during expedition TN401. We extend the same to the pilots and technical teams of ROVs *Victor 6000, Hyper-Dolphin, KM-ROV, Haima2, Isis, Ropos, SuBastian*, and *Jason II* as well as the HOVs *Nautile, Shinkai 6500, Pisces V*, and *Alvin* during those expeditions. We gratefully acknowledge the cruise PIs for leading the expeditions: François Lallier, Sorbonne Université (BIOBAZ); Didier Jollivet and Stéphane Hourdez, CNRS (CHUBACARC); Yves Fouquet (FUTUNA3); Cécile Cathalot and Ewan Pelleter, Ifremer (HERMINE2); Marjolaine Matabos and Jozée Sarrazin, Ifremer (MOMARSAT21); Karine Olu, Ifremer (WACS); Ken Takai, JAMSTEC (KY14-01, YK19-10, YK16-E02, YK22-05, KM23-E05); Masahiro Yamamoto, JAMSTEC (NT15-02, YK23-06); Shigeaki Kojima, AORI, the University of Tokyo (YK10-11); Takafumi Kasaya, JAMSTEC (NT15-02), Tatsuo Nozaki, Waseda University (KS-21-20); Hironori Komatsu, National Museum of Nature and Science (KS-22-2); Kenichiro Tani, National Museum of Nature and Science, Japan (YK23-16S); Jun Tao, Guangzhou Marine Geological Survey (HYDZ6-2020-05); Gary Massoth and Bob Embley, NOAA (KOK0505, KOK0506, KOK0507); Malcolm Clark, NIWA (TAN1206); Laura Wallace, New Zealand Institute of Geological and Nuclear Sciences (TAN2102); John W. Jamieson, Memorial University of Newfoundland (FKt231024) and Erik Cordes, Temple University (AT42-03). Biological sampling within the Commonwealth of the Northern Mariana Islands was carried out under the Marine Scientific Research permit number MSR U2022-047 from the United States government. This is a GACHINKO cruise Episode I (YK19-10) and Episode III (YK22-05) output. We are indebted to the Kingdom of Tonga and the government of Papua New Guinea for permitting access to their national waters and resources. We obtained the agreement to sample in Wallis et Futuna waters from the Haut Commissariat à la République in New Caledonia and the Préfecture in Wallis and Futuna. We thank Kareen Schnabel (NIWA) for providing samples from the Kermadec Arc and Hikurangi Margin, Aotearoa/New Zealand, Jianwen Qiu for providing samples from methane seep of the South China Sea and Hidetaka Nomaki (JAMSTEC) for providing facilities and discussions that helped in drafting this manuscript. We also thank Nicolas Gayet (UMR BEEP; Ifremer) and Katsuyuki Uematsu (Marine Work, Japan) for their work in preparing samples for scanning electron microscopy.

