## Supplementary Figures for "Evolutionary convergence and trophic diversity in hot vent and cold seep shrimps showcase a continuum of symbiosis"

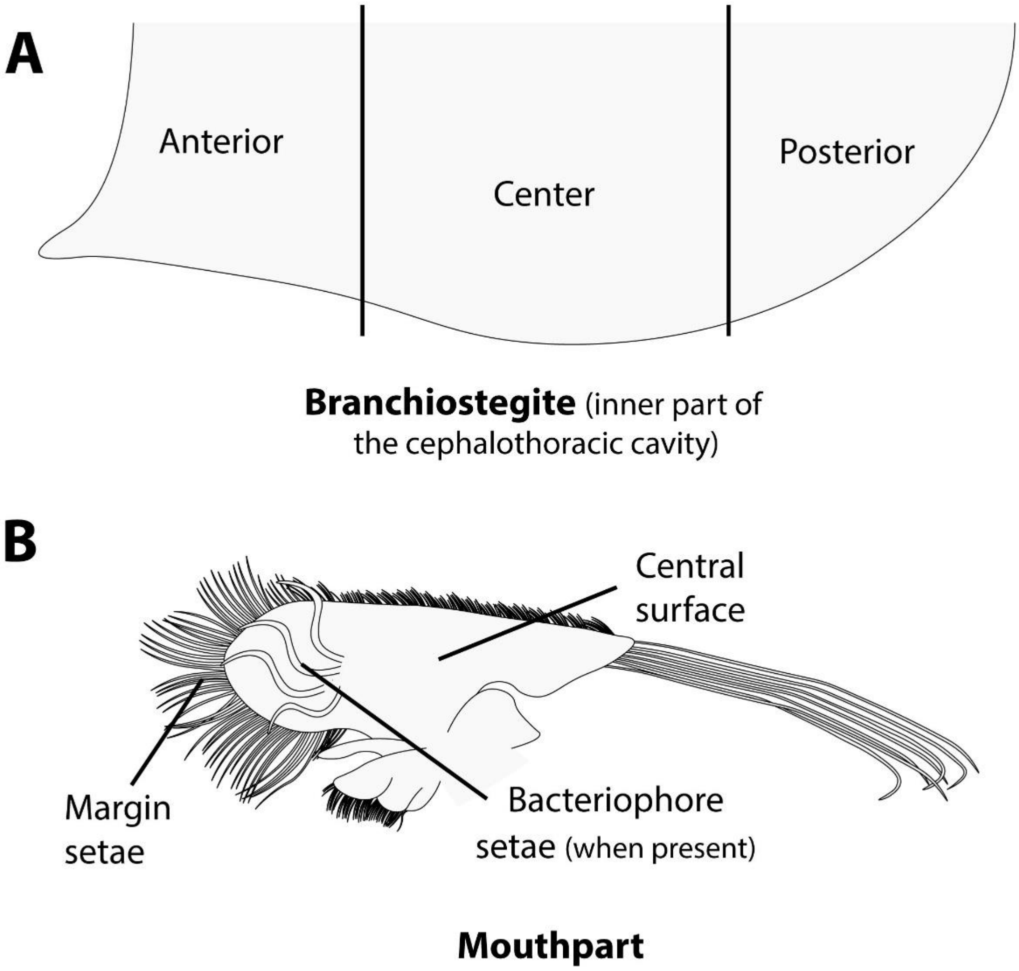


**Figure S1.** Schematic summarizing the method used to define bacterial colonisation in six distinct areas of the branchiostegite (anterior, center, posterior) or features of the mouthparts (central surface, margin setae, bacteriophore setae). For each area/feature, a score from 0 to 4 was attributed according to the following criteria, 0: no visible bacterial colonization; 1: a single layer of rod-shaped bacterial mats; 2: colonization by rod-shaped bacteria and sparsely distributed filamentous bacteria; 3: localized spots colonized by filamentous bacteria; 4: extremely dense colonization by filamentous bacteria, the surface of shrimp cuticle barely visible.

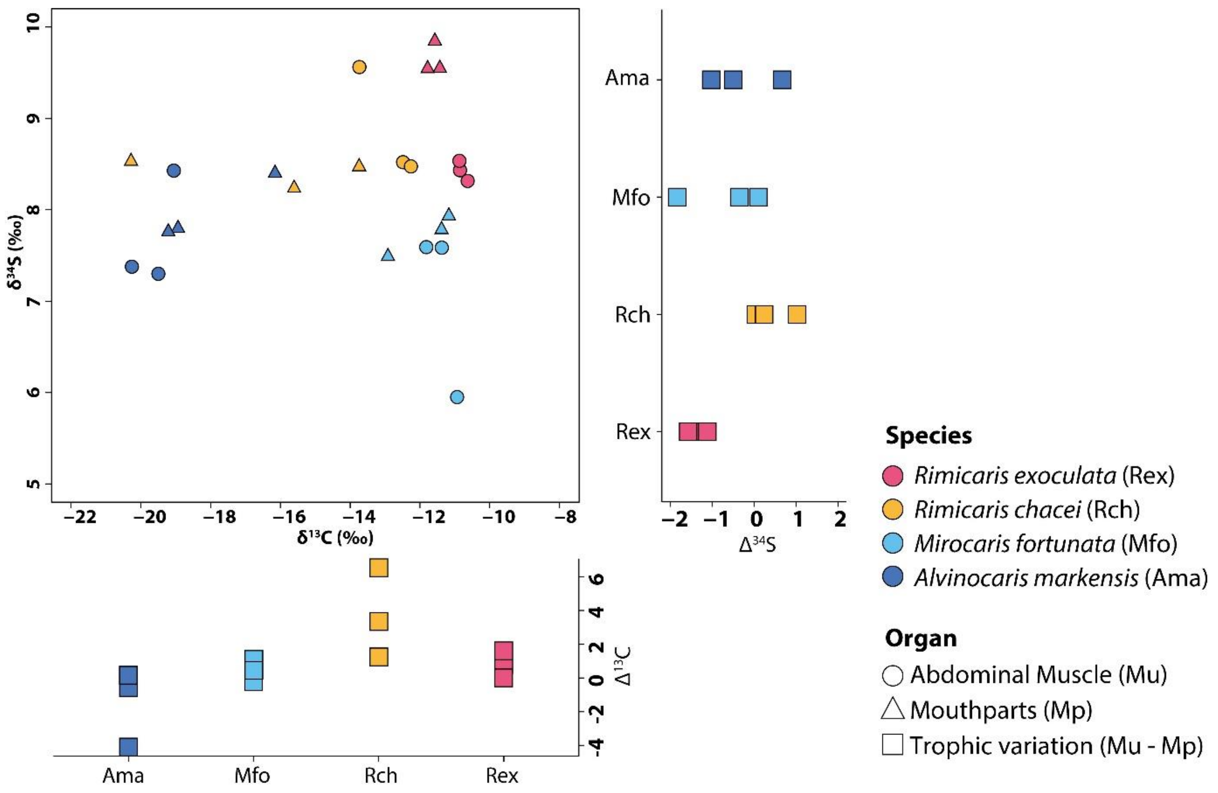


**Figure S2.** Carbon and sulfur isotopic ratios of abdominal muscles and mouthparts from alvinocaridid shrimps at the Puy des Folles vent field. $\Delta^{13}C$ and $\Delta^{34}S$ display variations of carbon and sulfur ratios between muscles and mouthpart tissues of each individual.

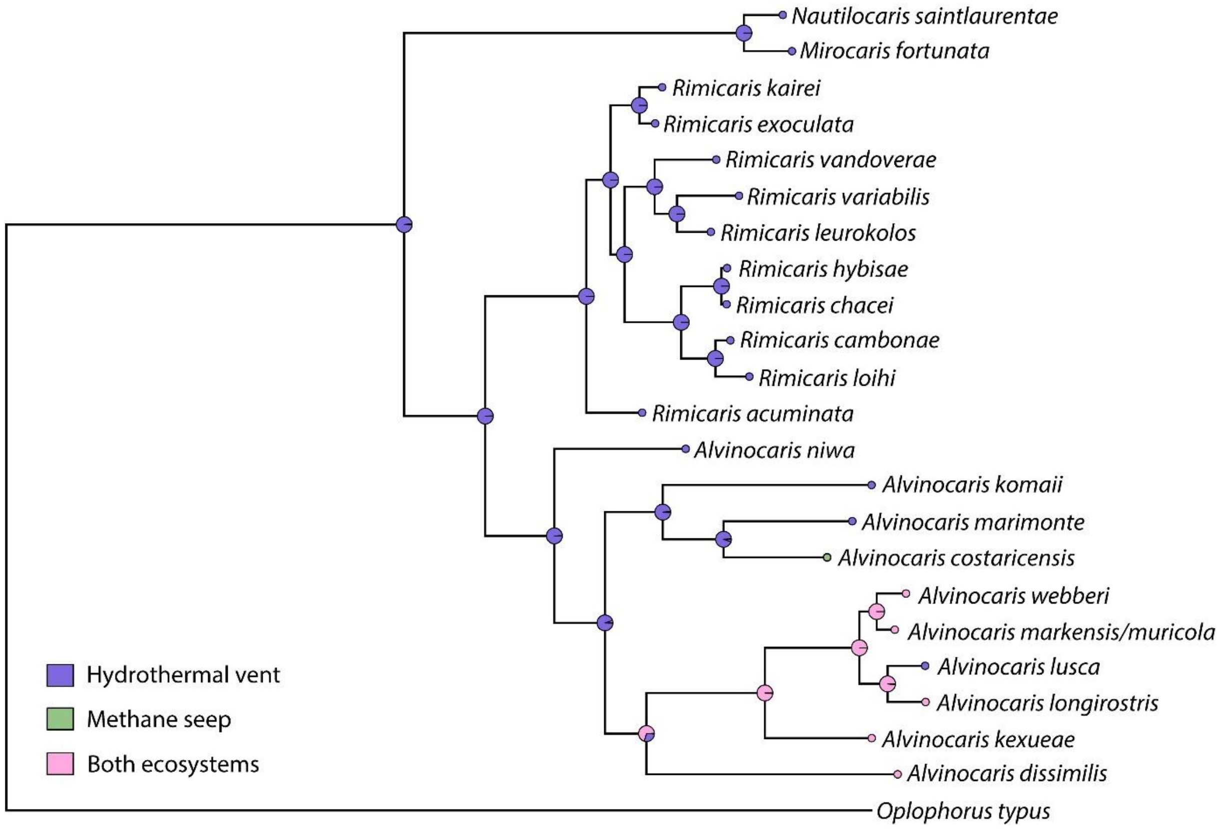


**Figure S3.** Phylogenetic tree of Alvinocarididae based on Bayesian inference (GTR + G + I model) generated from a 2177 bp concatenated alignment comprising partial fragments of the 18S, 16S, and COI genes. Pie charts represent ancestral state displaying posterior probabilities of each node being in each type of ecosystems.

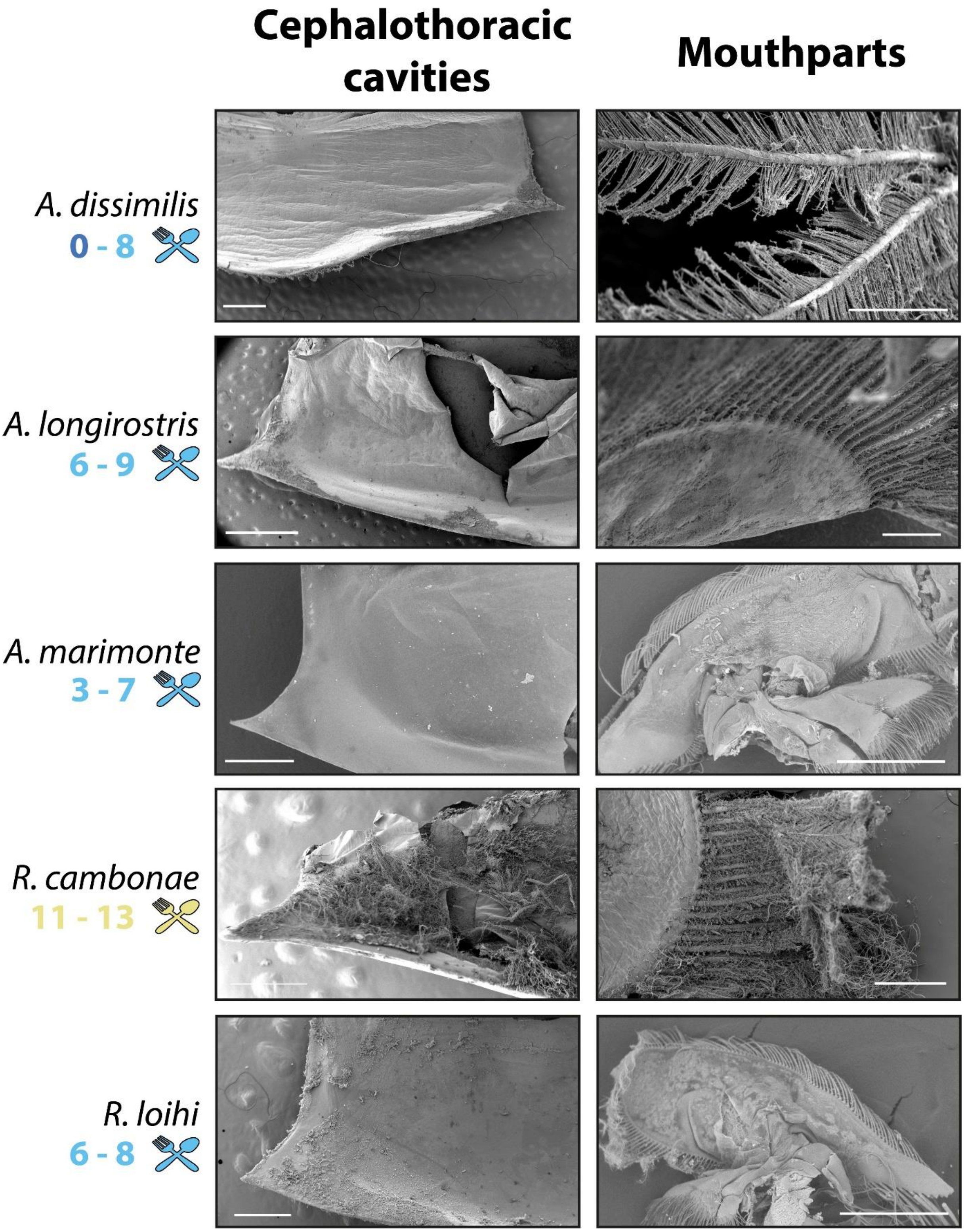


**Figure S4.** Additional examples of SEM observations for the cephalothoracic cavities and mouthparts for alvinocaridid species from the NW Pacific not displayed in Figure 2. Scale bars from top to bottom for cephalothoracic cavities: *A. dissimilis* (1 mm); *A. longirostris* (2 mm); *A. marimonte* (1 mm); *R. cambonae* (500 µm); *R. loihi* (500 µm). Scale bars from top to bottom for mouthparts: *A. dissimilis* (100 µm); *A. longirostris* (250 µm); *A. marimonte* (1 mm); *R. cambonae* (250 µm); *R. loihi* (1 mm).

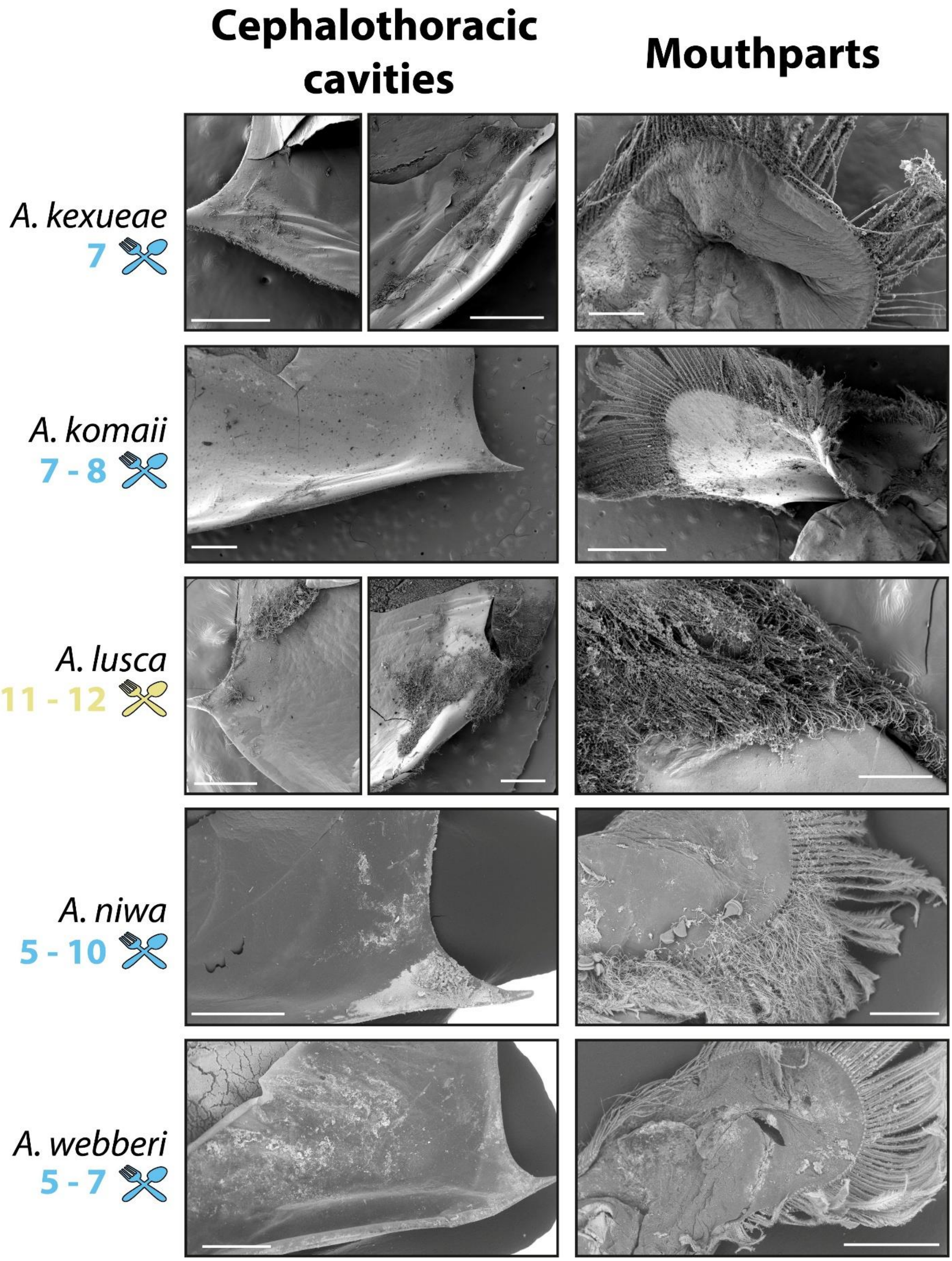


**Figure S5.** Additional examples of SEM observations for the cephalothoracic cavities and mouthparts for alvinocaridid species from the East & SW Pacific not displayed in Figure 2. Scale bars from top to bottom for cephalothoracic cavities: *A. kexueae* (anterior & posterior: 500 µm); *A. komaii* (1 mm); *A. lusca* (anterior & posterior: 500 µm); *A. niwa* (1 mm); *A. webberi* (1 mm). Scale bars from top to bottom for mouthparts: *A. kexueae* (500 µm); *A. komaii* (1 mm); *A. lusca* (500 µm); *A. niwa* (500 µm); *A. webberi* (1 mm).
